# Characterization of neutralizing versus binding antibodies and memory B cells in COVID-19 recovered individuals from India

**DOI:** 10.1101/2020.08.31.276675

**Authors:** Kaustuv Nayak, Kamalvishnu Gottimukkala, Sanjeev Kumar, Elluri Seetharami Reddy, Venkata Viswanadh Edara, Robert Kauffman, Katharine Floyd, Grace Mantus, Deepali Savargaonkar, Pawan Kumar Goel, Satyam Arora, Manju Rahi, Carl W Davis, Susanne Linderman, Jens Wrammert, Mehul S Suthar, Rafi Ahmed, Amit Sharma, Kaja Murali-Krishna, Anmol Chandele

**Affiliations:** ICGEB-Emory Vaccine Center, International Center for Genetic Engineering and Biotechnology, Aruna Asaf Ali Marg, New Delhi, India; Kasuma School of Biological Sciences, Indian Institute of Technology, New Delhi, India; Emory Vaccine Center, Emory University, Atlanta, GA 30322,USA; Department of Pediatrics, Emory University School of Medicine, Emory University, Atlanta, GA 30322, USA; Yerkes National Primate Research Center, Atlanta, GA 30329, USA; ICMR-National Institute of Malaria Research, Dwarka, New Delhi, India; Shaheed Hasan Khan Mewat Government Medical College, Nalhar, Mewat, Haryana, India; Department of Transfusion Medicine, Super Speciality Pediatric Hospital and Post Graduate Teaching Institute, Noida, UP, India; Division of Epidemiology and Communicable Diseases, Indian Council of Medical Research, New Delhi, India; Dept of Microbiology and Immunology, Emory University School of Medicine, Emory University, Atlanta, GA 30322, USA; Structural Parasitology Group, International Centre for Genetic Engineering and Biotechnology, Aruna Asaf Ali Marg, New Delhi, India

**Keywords:** SARS-CoV-2, COVID-19, Receptor Binding Domain, Neutralizing antibodies, Memory B cells, India

## Abstract

India is one of the countries most affected by the recent COVID-19 pandemic. Characterization of humoral responses to SARS-CoV-2 infection, including immunoglobulin isotype usage, neutralizing activity and memory B cell generation, is necessary to provide critical insights on the formation of immune memory in Indian subjects. In this study, we evaluated SARS-CoV-2 receptor-binding domain (RBD)-specific IgG, IgM, and IgA antibody responses, neutralization of live virus, and RBD-specific memory B cell responses in pre-pandemic healthy versus convalescent COVID-19 individuals from India. We observed substantial heterogeneity in the formation of humoral and B cell memory post COVID-19 recovery. While a vast majority (38/42, 90.47%) of COVID-19 recovered individuals developed SARS-CoV-2 RBD-specific IgG responses, only half of them had appreciable neutralizing antibody titers. RBD-specific IgG titers correlated with these neutralizing antibody titers as well as with RBD-specific memory B cell frequencies. In contrast, IgG titers measured against SARS-CoV-2 whole virus preparation, which includes responses to additional viral proteins besides RBD, did not show robust correlation. Our results suggest that assessing RBD-specific IgG titers can serve as a surrogate assay to determine the neutralizing antibody response. These observations have timely implications for identifying potential plasma therapy donors based on RBD-specific IgG in resource-limited settings where routine performance of neutralization assays remains a challenge.

**Importance:** Our study provides an understanding of SARS-CoV-2-specific neutralizing antibodies, binding antibodies and memory B cells in COVID-19 convalescent subjects from India. Our study highlights that PCR-confirmed convalescent COVID-19 individuals develop SARS-CoV-2 RBD-specific IgG antibodies, which correlate strongly with their neutralizing antibody titers. RBD-specific IgG titers, thus, can serve as a valuable surrogate measurement for neutralizing antibody responses. These finding have timely significance for selection of appropriate individuals as donors for plasma intervention strategies, as well as determining vaccine efficacy.

## Introduction

Severe acute respiratory syndrome coronavirus-2 (SARS-CoV-2), the virus responsible for the coronavirus disease 2019 (COVID-19) pandemic, emerged as a grave public health threat beginning in December 2019(1), paralyzing daily lives and causing economic downturns in many parts of the world. Currently, India is one of the countries most affected with more than 3 million COVID-19 confirmed cases and 60,000 associated deaths (2).

Intense efforts are underway to develop vaccines and antiviral therapeutics (3-11). These efforts require a detailed understanding of immune correlates of protection, formation of immune memory, and durability of these responses. Additionally, infusion of plasma derived from COVID-19 recovered individuals is also being explored as a treatment strategy (12-20). All these efforts require a detailed understanding of humoral immunity, immunoglobulin isotype usage and neutralizing activity following recovery from SARS-CoV-2 infection. Moreover, given that many of the SARS-CoV-2 neutralizing epitopes are located in the viral receptor binding domain (RBD) of the Spike (S) protein (21-29), it is important to evaluate the relationship between RBD-specific IgG titers and neutralizing antibody responses.

In this study, we evaluated IgG, IgA, IgM, neutralizing antibodies and memory B cell responses in PCR-confirmed COVID-19 convalescent subjects. Our results show that while a vast majority (38/42, 90.47%) of COVID-19 recovered individuals developed SARS-CoV-2 RBD-specific IgG responses, we were able to detect appreciable levels of neutralizing antibody responses in only half of the convalescent subjects. Neutralizing responses correlated closely with RBD-specific IgG titers, but weakly with IgG titers measured against crude virus concentrate using a commercial ELISA kit. Taken together, these findings suggest that despite significant inter-individual variation in the RBD-specific IgG titers and neutralizing antibodies, RBD-specific IgG titers can serve as a valuable and robust surrogate measurement for neutralizing antibody responses. These observations not only provide a glimpse of humoral immune responses in COVID-19 recovered individuals from India, but also have timely implications for identifying potential plasma therapy donors using on RBD-specific IgG ELISA’s in India where routine performance of neutralization assays remains a challenge.

## Methods

### Subject recruitment

COVID-19 recovered individuals were recruited at Shaheed Hasan Khan Mewati Government Medical College, Nuh, Haryana, India, Super Specialty Pediatric Hospital and Post Graduate Teaching Institute, Noida and ICMR-National Institute of Malaria Research, New Delhi. The Institutional ethical boards approved the study. Informed consent was obtained prior to inclusion in the study. All subjects (mean age 39.4 years, range 15 – 70 years) were SARS-CoV-2 PCR positive at the time of initial diagnosis, and were PCR negative when recruited for this study at 3.6 – 12 weeks post initial diagnosis **(Table 1)**. Samples collected from healthy adult blood bank donors in the year 2018 are included as pre-pandemic controls.

**Table 1.** COVID-19 recovered individuals characteristics (n=42)*

|  |  |
| --- | --- |
| Age in years Mean (Range) | 39.4 (15-70) |
| Males/Females | 38/4 |
| Days post PCR diagnosis Mean (Range) | 47.3 (25-84) |

\*COVID-19 recovered individuals were recruited at Shaheed Hasan Khan Mewati Government Medical College, Nuh, Haryana, India. Super Speciality Paediatric Hospital and Post Graduate Teaching Institute, Noida and ICMR-National Institute of Malaria Research, New Delhi. All subjects were SARS-CoV-2 PCR positive at the time of initial diagnosis and were PCR negative when recruited for this study at 4.8 – 11 weeks post initial diagnosis.

### SARS-CoV-2 *specific PCR*

SARS-CoV-2 specific rRT-PCR was performed as per the Indian government guidelines for COVID-19 diagnosis. Nasopharyngeal and throat swabs were collected in viral transport medium (VTM) (HiMedia, #AL 167)) and transported to the testing laboratory maintaining cold chain. All the samples were subjected to the first line screening assay or the ‘e’ gene assay as per the guidelines (30). Samples reactive by the first line assay were subjected to the RdRp gene assay (Invitrogen SuperScript™ III Platinum® One-Step Quantitative Kit (Cat. No.11732088). Samples reactive for both the genes were labeled positive, while samples reactive to ‘e’ gene only were considered indeterminate and were subjected to repeat sampling. The same protocol was used to verify that the subjects were PCR negative at the time of recruitment for this study.

### SARS-CoV-2 *RBD-specific direct ELISA*

Recombinant SARS-CoV-2 RDB gene was cloned, expressed, purified and standard direct ELISAs were performed as previously described (31). Briefly, purified RBD was coated on MaxiSorp plates (Thermo Fisher, #439454) at a concentration of 1 ug/mL in 100 uL phosphate-buffered saline (PBS) at 4°C overnight. The plates were washed extensively with PBS containing 0.05% Tween-20. Three-fold serially diluted plasma samples were added to the plates and incubated at room temperature for 1hr. After incubation, the plates were washed and the SARS-CoV-2 RBD specific IgG, IgM, IgA signals were detected by incubating with horseradish peroxidase (HRP) conjugated - anti-human IgG (Jackson ImmunoResearch Labs, #109-036-098), IgM (Jackson ImmunoResearch Labs, #109-036-129), or IgA (Jackson ImmunoResearch Labs, #109-036-011). Plates were then washed thoroughly and developed with o-phenylenediamine (OPD) substrate (Sigma, #P8787) in 0.05M phosphate-citrate buffer (Sigma, #P4809) pH 5.0, containing with 0.012% hydrogen peroxide (Fisher Scientific, #18755) just before use. Absorbance was measured at 490 nm.

### *Enumeration of* SARS-CoV-2 *RBD-specific memory B cells*

Purified RBD protein (100 ug) was labeled with Alexa Fluor 488 using microscale protein labeling kit (Life Technologies, #A30006) as per manufacturer’s protocol. PBMC’s were stained with RBD-Alexa Fluor 488 for 1 hour at 4°C, followed by washing with PBS containing 0.25% FBS, and incubation with efluor780 Fixable Viability (Live Dead) dye (Life Technologies, #65-0865-14) and anti-human CD3, CD19, CD27, CD38 and IgD antibodies (BD Biosciences) for 30 minutes. Cells were washed twice with FACS buffer and acquired on BD LSR Fortessa X20. Data was analyzed using FlowJo software 10. SARS-CoV-2 RBD-specific memory B cells were identified in cells positive for CD19, CD20, CD27 that were negative for IgD and CD3.

### IgG ELISA for SARS-CoV-2 whole virus preparation

SARS-CoV-2 antigen specific IgG was detected using a commercially available assay (COVID-Kavach ELISA tests kit, Zydus diagnostics), which measures responses to antigen concentrated from gamma-irradiated SARS-CoV-2-infected tissue culture fluid as per the manufacturer’s instructions (32, 33).

### SARS-CoV-2 neutralization assay

Neutralization titers to SARS-CoV-2 were determined as previously described (31). Briefly infectious clone of the full-length mNeonGreen SARS-CoV-2 (2019-nCoV/USA_WA1/2020) was used to test heat-inactivated COVID-19 convalescent samples and healthy donor samples (pre-pandemic). Heat-inactivated serum was serially diluted three-fold in duplicate starting at a 1:20 dilution in a 96-well round-bottom plate and incubated between 750 FFU of ic-SARS-CoV-2-mNG for 1 h at 37°C. This antibody-virus mixture was transferred into the wells of a 96-well plate that had been seeded with Vero-E6 cells the previous day at a concentration of 2.5× 10^4^ cells/well. After 1 hour, the antibody-virus inoculum was removed and 0.85% methylcellulose in 2% FBS containing DMEM was overlaid onto the cell monolayer. Cells were incubated at 37°C for 24 hours. Cells were washed three times with 1XPBS (Corning Cellgro) and fixed with 125 µl of 2% paraformaldehyde in PBS (Electron Microscopy Sciences) for 30 minutes. Following fixation, plates were washed twice with 1x PBS and imaged on an ELISPOT reader (CTL Analyzer). Foci were counted using Viridot (34) (counted first under the “green light” setting followed by background subtraction under the “red light” setting). FRNT-mNG_50_ titers were calculated by non-linear regression analysis using the 4PL sigmoidal dose curve equation on Prism 8 (Graphpad Software). Neutralization titers were calculated as 100% x [1-(average foci in duplicate wells incubated with the specimen) ÷ (average number of foci in the duplicate wells incubated at the highest dilution of the respective specimen).

### Statistical analysis

Statistical analysis was performed using GraphPad prism 8.0 software. Non-parametric t test (Mann-Whitney) was used to calculate the differences between groups. Non-parametric Spearman’s correlation coefficient (r) was used to calculate correlation between groups. A *p* value of ≤0.05 was considered as significant.

## Results

### SARS-CoV-2 RBD-specific humoral immunity in COVID-19 recovered individuals

The demographic profile of COVID-19 recovered individuals recruited for this study is shown in **Table 1**. All subjects were at least 3.6 weeks past their initial SARS-CoV-2 positive diagnosis. RBD-specific ELISA curves for IgG, IgA and IgM at different dilutions of plasma in pre-pandemic healthy versus COVID-19 recovered individuals are shown in **Figure 1**. RBD-specific responses were highly elevated in COVID-19 recovered individuals as compared to pre-pandemic healthy controls (**Figure 1A,B,C, left versus middle panels)**. Titers of IgG, IgA and IgM in the COVID-19 recovered individuals showed substantial inter-individual variation **(Figure 1 A, B, C, right panel)** - with IgG endpoint titers ranging from below detection to 24484 (2000±619); IgA titers from below detection to 5686 (386±136) and IgM titers from below detection to 2958 (515±90). Four individuals had undetectable RBD-specific IgG and IgA titers. One of these individuals was also below detection for IgM **(Table 2)**. Inter-individual heterogeneity was not related to the age of the individuals **(Figure 2A)** or the number of days that elapsed between PCR confirmation of infection and sample collection **(Figure 2B)**.

**Table 2.**
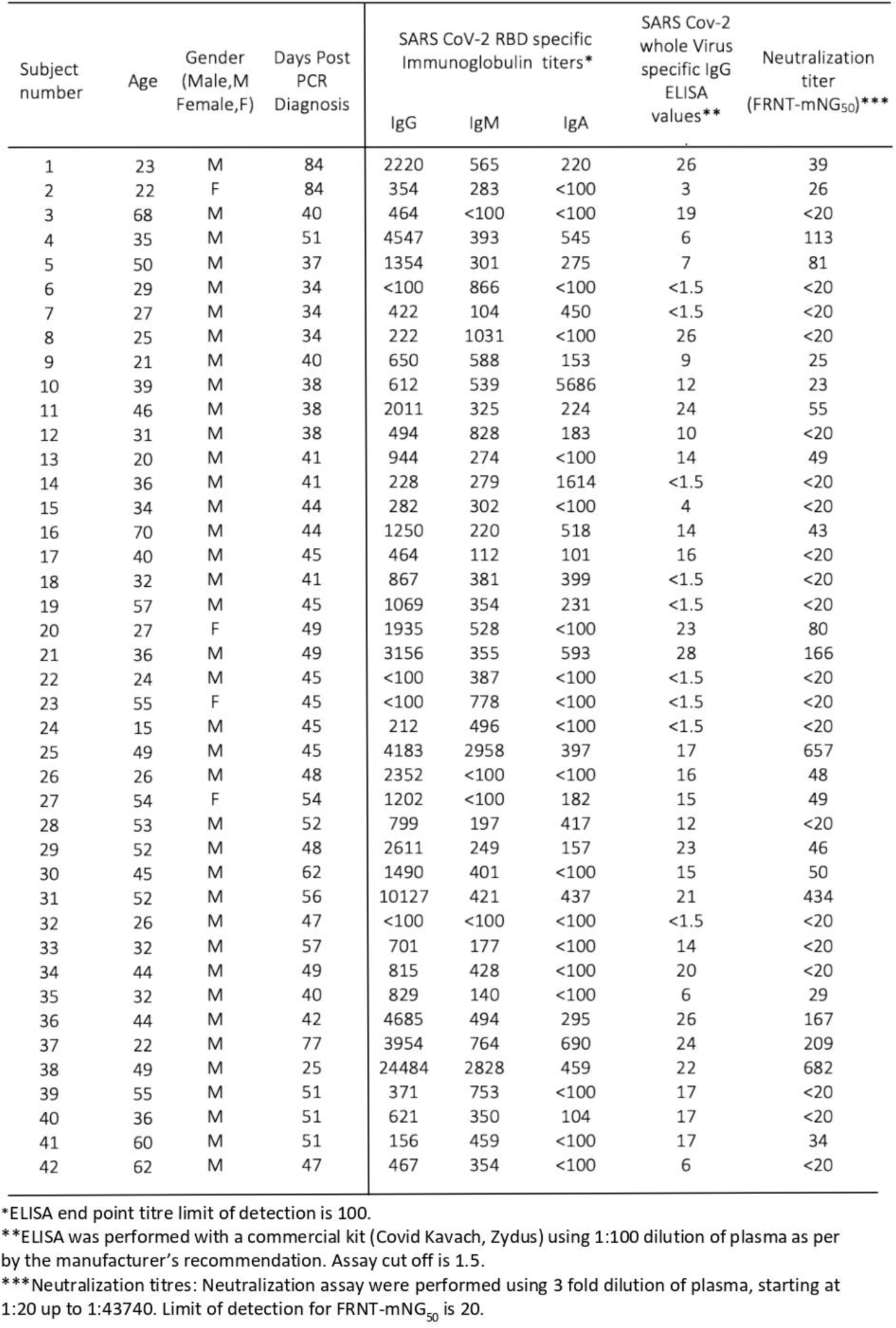
Individual characteristics of the COVID-19 recovered subjects.

| Subject number | Age | Gender (Male,M Female,F) | Days Post PCR Diagnosis | SARS CoV-2 RBD specific Immunoglobulin titers* |  |  | SARS Cov-2 whole Virus specific IgG ELISA values** | Neutralization titer (FRNT-mNG<sub>50</sub>)*** |
| --- | --- | --- | --- | --- | --- | --- | --- | --- |
|  |  |  |  | IgG | IgM | IgA |  |  |
| 1 | 23 | M | 84 | 2220 | 565 | 220 | 26 | 39 |
| 2 | 22 | F | 84 | 354 | 283 | <100 | 3 | 26 |
| 3 | 68 | M | 40 | 464 | <100 | <100 | 19 | <20 |
| 4 | 35 | M | 51 | 4547 | 393 | 545 | 6 | 113 |
| 5 | 50 | M | 37 | 1354 | 301 | 275 | 7 | 81 |
| 6 | 29 | M | 34 | <100 | 866 | <100 | <1.5 | <20 |
| 7 | 27 | M | 34 | 422 | 104 | 450 | <1.5 | <20 |
| 8 | 25 | M | 34 | 222 | 1031 | <100 | 26 | <20 |
| 9 | 21 | M | 40 | 650 | 588 | 153 | 9 | 25 |
| 10 | 39 | M | 38 | 612 | 539 | 5686 | 12 | 23 |
| 11 | 46 | M | 38 | 2011 | 325 | 224 | 24 | 55 |
| 12 | 31 | M | 38 | 494 | 828 | 183 | 10 | <20 |
| 13 | 20 | M | 41 | 944 | 274 | <100 | 14 | 49 |
| 14 | 36 | M | 41 | 228 | 279 | 1614 | <1.5 | <20 |
| 15 | 34 | M | 44 | 282 | 302 | <100 | 4 | <20 |
| 16 | 70 | M | 44 | 1250 | 220 | 518 | 14 | 43 |
| 17 | 40 | M | 45 | 464 | 112 | 101 | 16 | <20 |
| 18 | 32 | M | 41 | 867 | 381 | 399 | <1.5 | <20 |
| 19 | 57 | M | 45 | 1069 | 354 | 231 | <1.5 | <20 |
| 20 | 27 | F | 49 | 1935 | 528 | <100 | 23 | 80 |
| 21 | 36 | M | 49 | 3156 | 355 | 593 | 28 | 166 |
| 22 | 24 | M | 45 | <100 | 387 | <100 | <1.5 | <20 |
| 23 | 55 | F | 45 | <100 | 778 | <100 | <1.5 | <20 |
| 24 | 15 | M | 45 | 212 | 496 | <100 | <1.5 | <20 |
| 25 | 49 | M | 45 | 4183 | 2958 | 397 | 17 | 657 |
| 26 | 26 | M | 48 | 2352 | <100 | <100 | 16 | 48 |
| 27 | 54 | F | 54 | 1202 | <100 | 182 | 15 | 49 |
| 28 | 53 | M | 52 | 799 | 197 | 417 | 12 | <20 |
| 29 | 52 | M | 48 | 2611 | 249 | 157 | 23 | 46 |
| 30 | 45 | M | 62 | 1490 | 401 | <100 | 15 | 50 |
| 31 | 52 | M | 56 | 10127 | 421 | 437 | 21 | 434 |
| 32 | 26 | M | 47 | <100 | <100 | <100 | <1.5 | <20 |
| 33 | 32 | M | 57 | 701 | 177 | <100 | 14 | <20 |
| 34 | 44 | M | 49 | 815 | 428 | <100 | 20 | <20 |
| 35 | 32 | M | 40 | 829 | 140 | <100 | 6 | 29 |
| 36 | 44 | M | 42 | 4685 | 494 | 295 | 26 | 167 |
| 37 | 22 | M | 77 | 3954 | 764 | 690 | 24 | 209 |
| 38 | 49 | M | 25 | 24484 | 2828 | 459 | 22 | 682 |
| 39 | 55 | M | 51 | 371 | 753 | <100 | 17 | <20 |
| 40 | 36 | M | 51 | 621 | 350 | 104 | 17 | <20 |
| 41 | 60 | M | 51 | 156 | 459 | <100 | 17 | 34 |
| 42 | 62 | M | 47 | 467 | 354 | <100 | 6 | <20 |

\*ELISA end point titre limit of detection is 100.
\*\*ELISA was performed with a commercial kit (Covid Kavach, Zydus) using 1:100 dilution of plasma as per by the manufacturer's recommendation. Assay cut off is 1.5.
\*\*\*Neutralization titres: Neutralization assay were performed using 3 fold dilution of plasma, starting at 1:20 up to 1:43740. Limit of detection for FRNT-mNG<sub>50</sub> is 20.

**Figure 1:**
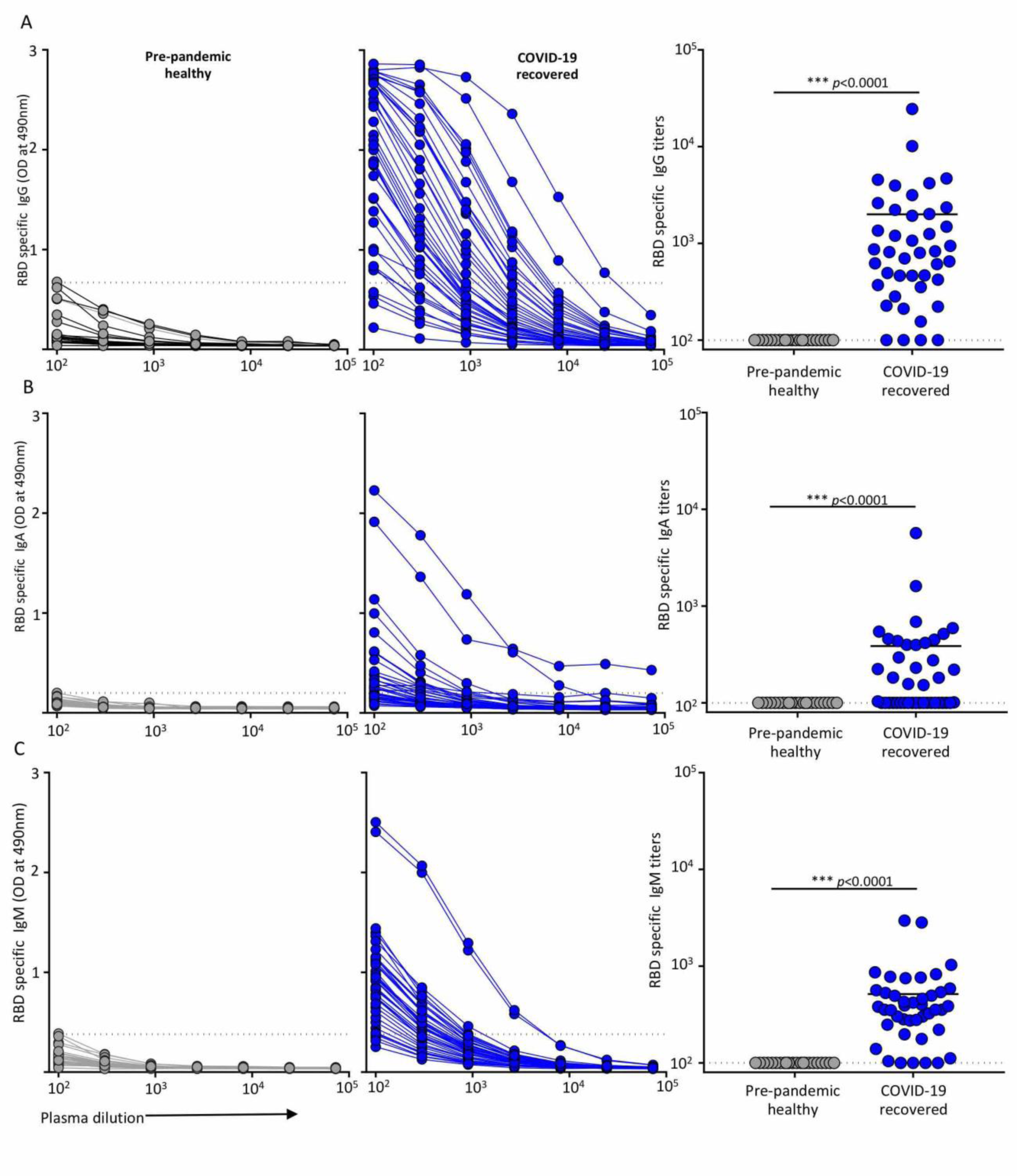
Evaluation of SARS-CoV-2 RBD specific IgG, IgA and IgM antibody responses. **(A)** RBD-specific IgG, (**B)**, RBD-specific IgA; (**C)**, RBD-specific IgM. Left, pre-pandemic healthy (n-22), middle COVID-19 recovered (n=42); right, endpoint titers. ELISA cutoff values are calculated using the average plus 3 standard deviations of the 22 healthy controls at 1:100 dilution (shown as a dotted line). The unpaired analysis was done using non-parametric Mann-Whitney-U test. *p* ≤ 0.05 was considered significant. Assay cutoff value is marked with dotted line.

**Figure 2.**
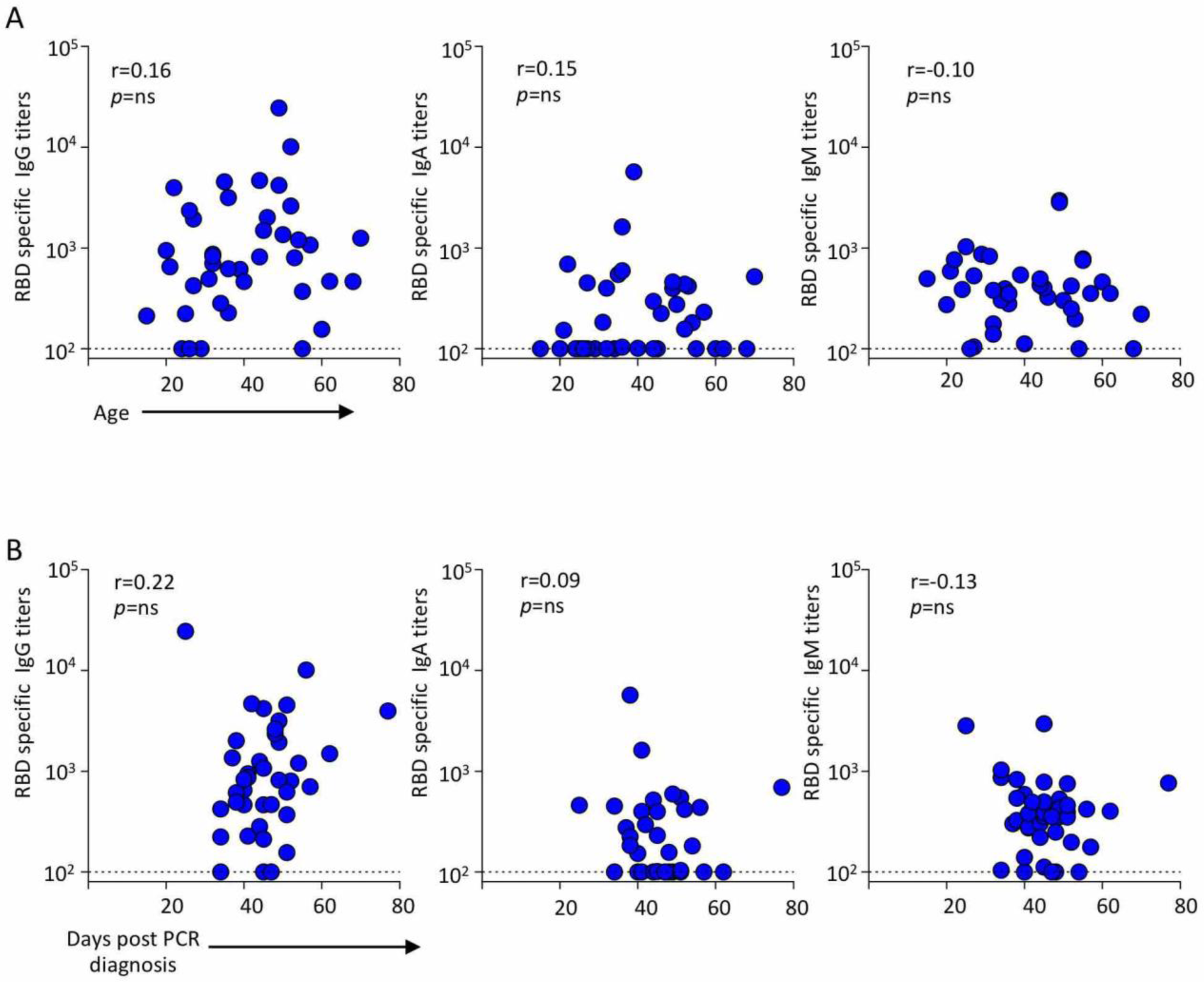
Correlation of age and day post initial diagnosis of COVID-19 recovered individuals with SARS-CoV-2 IgG, IgM and IgA titers. **(A)**. Age versus IgG (left, n=42), IgA (middle, n=42) or IgM (right, n=42) titers. (**B)**. Time post initial diagnosis versus IgG (left, n=42), IgA (middle, n=42) or IgM (right, n=42) titers. Correlations were calculated by Spearman’s correlation coefficient r. *p* ≤ 0.05 is considered significant. Note that none of the data sets above reached significant values of correlation.

### SARS-CoV-2 specific neutralizing titers in COVID-19 recovered individuals

To assess plasma neutralizing titers from COVID-19 convalescent individuals, we performed a live virus neutralization assay using a focus-reduction neutralization mNeonGreen (FRNT-mNG) assay (31). The neutralizing activity at different dilutions of plasma for pre-pandemic healthy individuals **(Figure 3A)** and COVID-19 recovered individuals is shown in (**Figure 3B). Figure 3C** shows FRNT-mNG_50_ titers calculated based on the plasma dilution that neutralized 50% of the virus. While all pre-pandemic healthy individuals had undetectable FRNT-mNG_50_ titers, only half of the COVID-19 recovered individuals showed 50% or more neutralization even at a 1:20 dilution of plasma. Similar to RBD-specific IgG titers, the FRNT-mNG_50_ titers were heterogeneous with the latter reaching titers as high as 682 **(Figure 3C)**.

**Figure 3.**
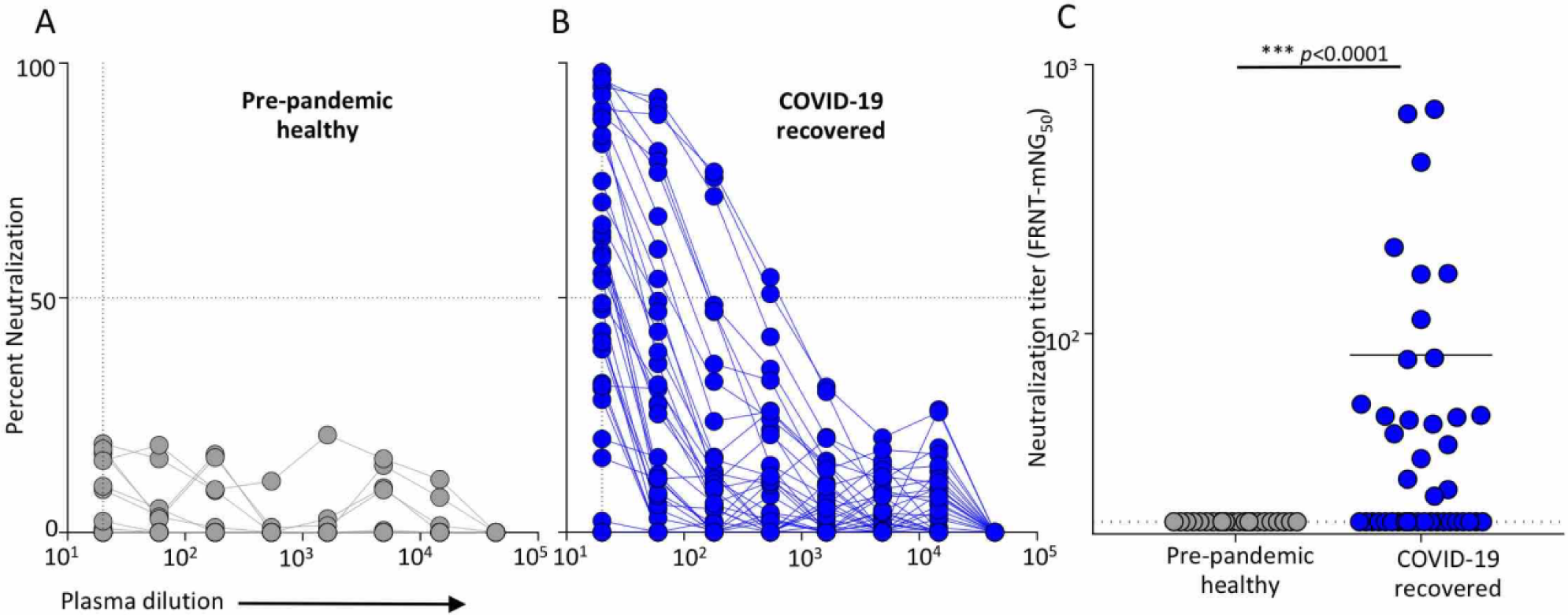
Evaluation of SARS-CoV-2 neutralizing antibodies in COVID-19 recovered individuals. SARS-CoV-2 neutralizing activity at indicated dilutions of plasma is shown in pre-pandemic healthy (n=22, in grey) **(A)** and in COVID-19 recovered individuals (n=42, in blue) **(B)**. Dotted line represents the plasma dilution that leads to 50% neutralization. **(C)** Scatter plot shows neutralization titers (FRNT-mNG_50_) in pre-pandemic healthy (n=22) and COVID-19 recovered (n=42) individuals. The unpaired analysis was done using non-parametric Mann-Whitney-U test. *p* ≤ 0.05 was considered significant. Limit of detection is marked with a dotted line.

Previous studies in other viral infections have shown that all three antibody isotypes (IgG, IgA and IgM) can potentially neutralize (35-39). We next determined if any correlation exists between SARS-CoV-2 neutralizing titers and RBD-specific IgG, IgA, IgM binding antibody titers. We observed a positive correlation (r=0.83; p<0.001) between SARS-CoV-2 neutralizing titers and RBD-specific IgG titers **(Figure 4, left graph)** but not with IgA **(Figure 4, middle graph)** or IgM titers **(Figure 4, right graph)**.

**Figure 4.**
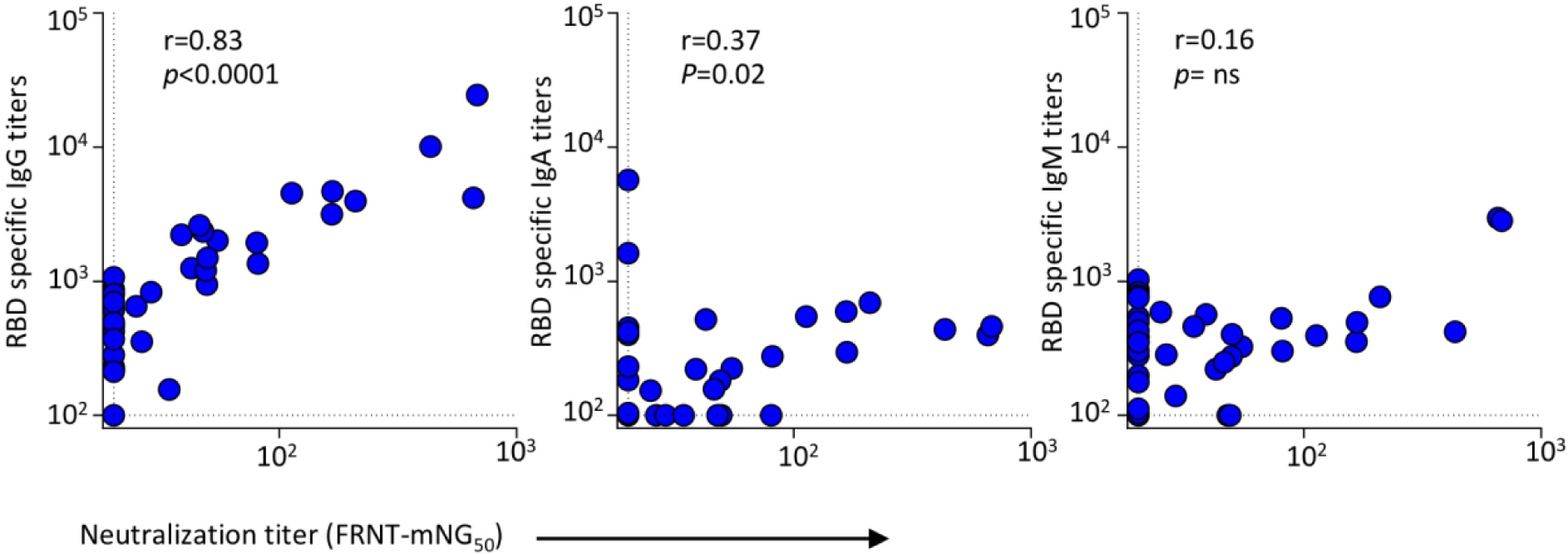
Correlation analysis of SARS-CoV-2-specific antibody responses versus neutralization titers. Correlation analysis shows FRNT-mNG_50_ titers (x-axis) versus RBD-specific IgG (Left), IgA (middle) and IgM (right) titers on y-axis in COVID-19 recovered individuals (n=42, blue dots). Correlation analysis was performed by log transformation of the endpoint ELISA titers followed by linear regression analysis. Correlations were calculated by Spearman’s correlation coefficient r. *p* ≤ 0.05 was considered significant. Dotted line on x-axis and y-axis indicate limit of detection.

Plasma infusion therapy has recently been started in India as an intervention therapy for COVID-19. For this, plasma donors are being typically identified by the presence of IgG to SARS-CoV-2 by commercial ELISA tests (40). One of these tests detects IgG towards viral antigens concentrated from gamma-irradiated SARS-CoV-2-infected tissue culture fluid (32, 33). It was therefore of interest to examine the correlation between neutralization titers and IgG responses measured using this test. We observed that, of the 42 COVID-19 recovered individuals tested, 33 were IgG positive whereas 9 were below the assay cut off **(Figure 5A)**. Of the 9 individuals that were below cut off, 4 also tested negative by the RBD-specific IgG ELISA **(Table 2)**. All of the samples from the pre-pandemic healthy individuals were below the limit of detection using both the ELISA methods. Most importantly, the IgG values obtained by whole virus-based ELISA did not show as robust a correlation (r=0.56) with neutralizing antibody titers **(Figure 5B)** as compared to those observed with RBD-specific IgG titers (r=0.83) **(Figure 4, left graph)**.

**Figure 5.**
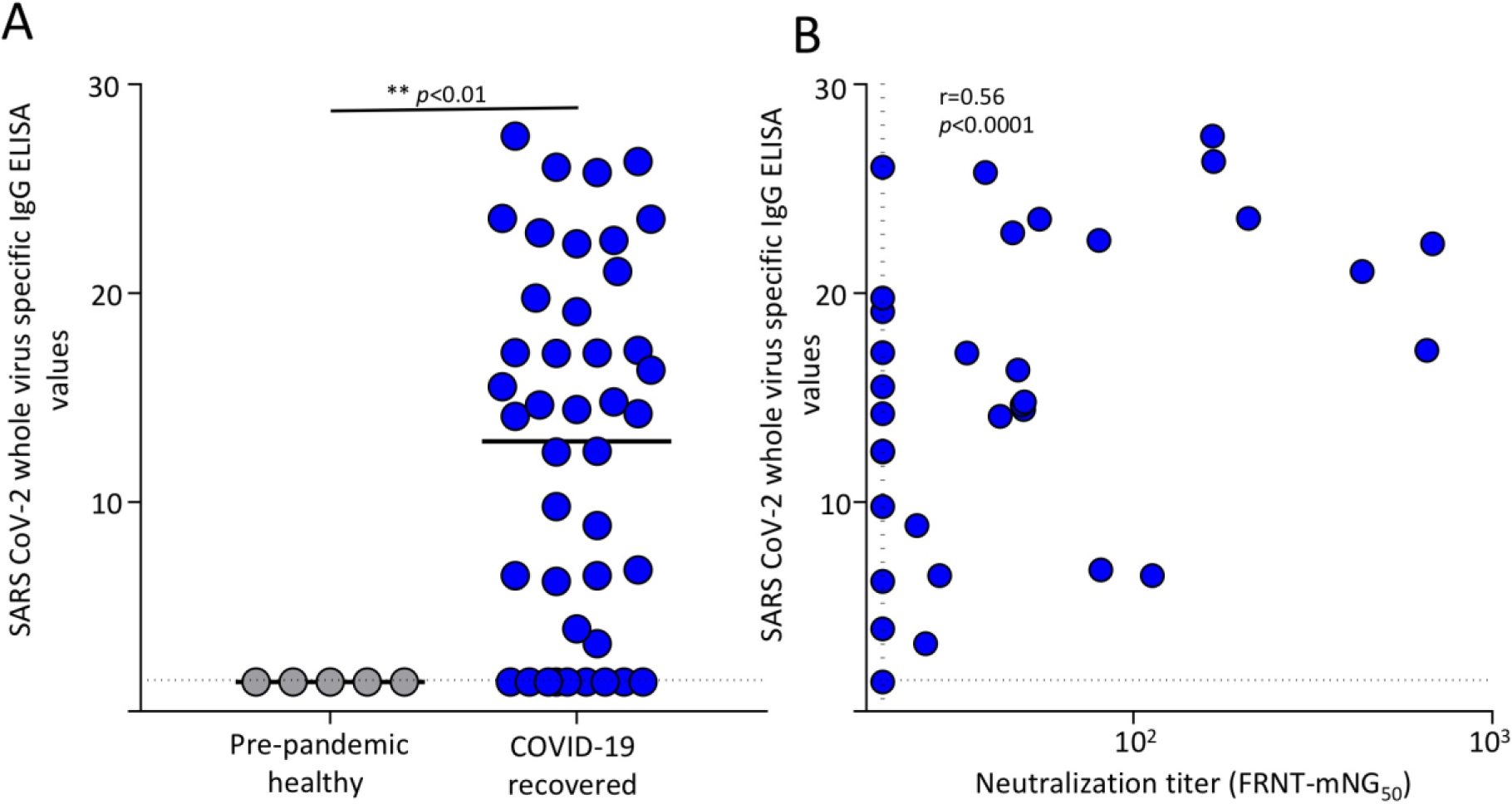
Correlation analysis of SARS-CoV-2 whole virus specific IgG versus neutralizing titers. **(A)**. Scatter plots shows SARS-CoV-2 whole virus specific IgG measured using measured using commercial kit (Zydus diagnosis, Covid Kavach) in pre-pandemic healthy (n=5) and COVID-19 recovered (n=42). The unpaired analysis was done using non-parametric Mann-Whitney-U test. *p* ≤ 0.05 was considered significant. **(B)**. Correlation analysis of SARS-CoV-2 whole virus antigen specific IgG ELISA kit values (y-axis) versus neutralizing titers (x-axis) in COVID-19 recovered individuals (n=42). Correlations were calculated by Spearman’s correlation coefficient r. p ≤ 0.05 was considered significant. Dotted line on x-axis indicate limit of detection and on y-axis assay cut off.

### Characterization of RBD-specific memory B cells in COVID-19 recovered individuals

While circulating neutralizing antibodies help prevent re-infection by viruses, memory B cells allow for rapid production of new antibodies in case of re-infection. To address whether the COVID-19 recovered individuals generated memory B cells, we enumerated RBD-specific memory B cells using fluorescently-conjugated RBD antigen. An example of the flow cytometric gating strategy and RBD staining among the gated memory B cells is shown in **Figure 6A and 6B. Figure 6C** shows the frequency of RBD-specific memory B cells in a subset of the individuals where sufficient PBMCs were available. Though we found that there was substantial inter-individual variation in the frequency of SARS-CoV-2 RBD-specific memory B cells, their frequencies modestly correlated with RBD-specific IgG titers.

**Figure 6.**
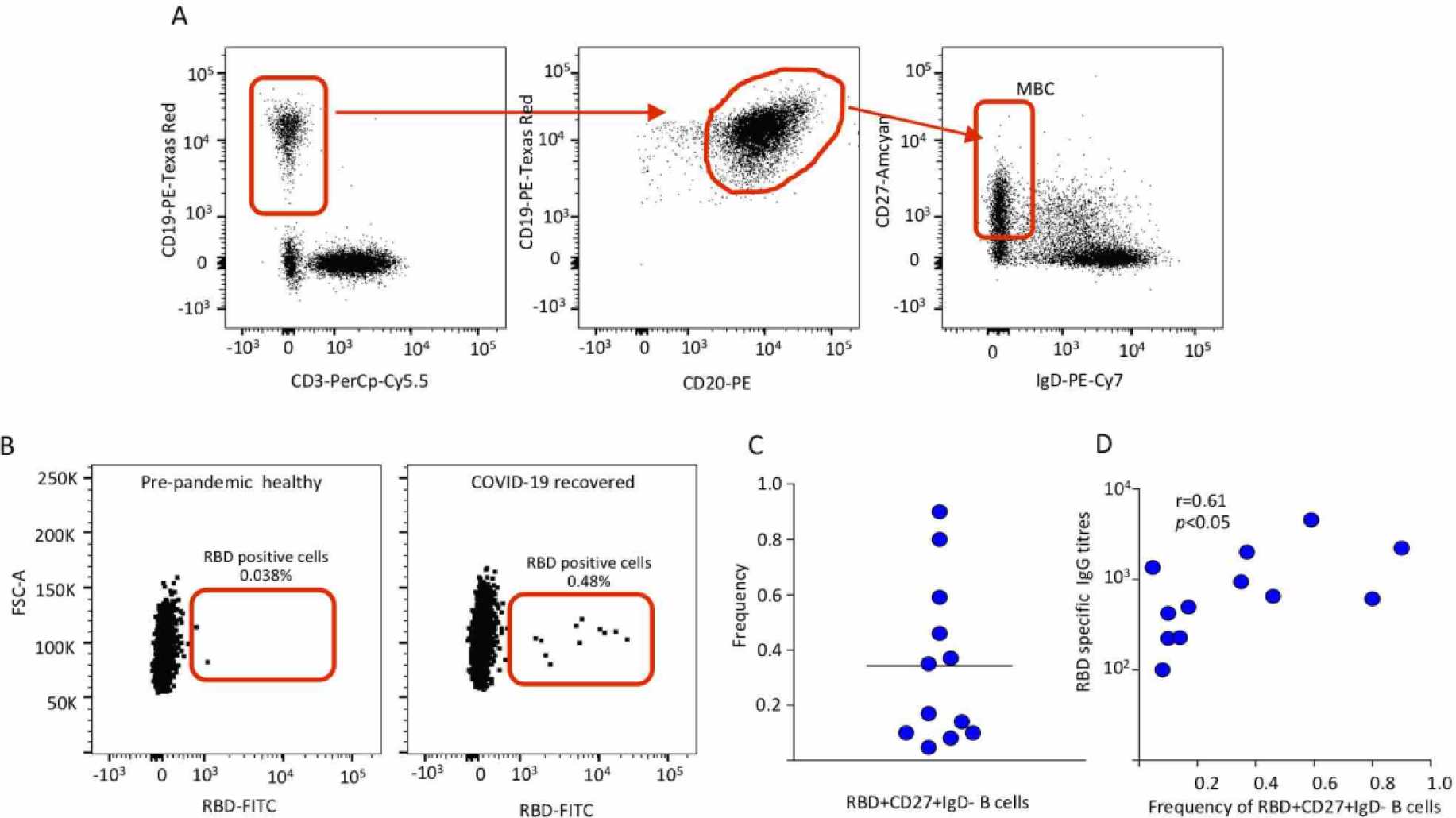
SARS-CoV-2 RBD-specific memory B cell analysis in COVID-19 recovered individuals. **(A)** Gating strategy used to identify memory B cells. **(B)** SARS-CoV-2 RBD-specific memory B cells on gated total memory B cells that were CD19 positive, CD20 high, IgD negative and CD27 high is shown. **(C)** Frequency of RBD-specific memory B cells of the total memory B cells in the COVID-19 recovered individuals (n= 13). **(D)** Correlation analysis shows frequency of RBD-specific memory B cells (x-axis) and the RBD-specific IgG titers (y-axis) in COVID-19 recovered individuals.

## Discussion

Our study provides a detailed understanding of humoral immunity and memory B cells in COVID-19 recovered individuals from India. We examined SARS-CoV-2 neutralizing antibodies, IgG, IgM, IgA and memory B cells in pre-pandemic healthy versus COVID-19 recovered individuals and further evaluated inter-individual variation and relation among these.

Our correlative analysis of RBD-specific IgG binding titers with neutralizing antibody titers and memory B cells has important implications for not only identifying potential donors for plasma therapy but also for understanding humoral and cellular memory post COVID-19. Though current plasma therapy guidelines in India do not consider neutralizing antibody titers, United States Food and Drug Administration (FDA) guidelines recommend, when available, a neutralizing titer of 1:160 or 1:80 to be used for identifying potential plasma donors (41). Our correlation analysis shows that RBD-specific titers of more than 3668 can provide a suitable surrogate for identifying the individuals with neutralizing titers of above 1:160 and RBD-specific IgG titers 1926 for neutralizing titers of 1:80. Though larger scale studies are needed to establish robustness, these observations have timely implications to identify potential plasma therapy donors.

Our study raises important questions on formation of protective immune memory after recovering from COVID-19. We found that nearly half of the COVID-19 recovered individuals did not induce 50% neutralizing titers even at 1:20 dilution of plasma. This raises the question of whether these individuals with low neutralizing antibodies also differ in formation of cellular immune memory. Our data show that individuals with low neutralizing antibodies indeed had lower memory B cells. Given that T cells may also contribute to COVID-19 protection, studies are needed to understand whether these individuals may also differ in the generation of memory CD8 and CD4 T cells (42-44).

The reason why only half of the COVID-19 recovered individuals developed appreciable levels of neutralizing antibody titers requires further investigation. This may be related to inter-individual differences in human immune responses associated with the expected heterogeneity in initial viral inoculum(45), initial viral loads (46-48), incubation period (49), host genetic factors (50-52) and disease severity (53, 54). This is consistent with previous studies that show relatively higher neutralizing antibodies in COVID-19 hospitalized patients during the acute febrile phase, or in recovered individuals that were previously hospitalized with severe COVID-19 disease (53, 54). It is noteworthy that the COVID-19 recovered individuals from our study had mild to moderate symptoms during the initial diagnosis. In light of these studies, our findings warrant future studies to seek an understanding of whether the individuals that have generated low or no neutralizing antibodies, IgG titers or memory B cells past recovery will be protected if they were re-exposed to SARS-CoV-2 or a related virus.

## Acknowledgements

This research was supported in part by Indian Council of Medical Research VIR/COVID-19/02/2020/ECD-1 (A.C, K.M). K.N, E.S.R. are supported through Dengue Translational Research Consortia BIRAC/NBM-PMU/EST-TRC_ICGEB/2018 (A.C.); K.G. is supported through DBT grant BT/PR30260/MED/15/194/2018 (A.C, K.M); S.K. is supported through DBT/Wellcome Trust India Alliance Early Career Fellowship grant IA/E/18/1/504307. We are thankful to Mr. Satendra Singh and Mr. Ajay Singh, ICGEB, New Delhi for technical support; Director, SSPH & PGTI, Noida and Director, SHKM Government Medical College, Nuh, Haryana for facilitating the study. Vineet Menachery and Pei-Yong Shi for providing the icSARS-CoV-2mNG for the neutralization assays.

## Author Contributions

Experimental work, data acquisition and analysis of data by K.N, K.G, S.K, E.S.R, V.V.E., K.F, R.K, S.L. C.D, J.W, M.S.S, and D.S. Clinical site coordination by D.S, P.K.G, S.A, A.S and M.R. Conceptualization and implementation by A.S, R.A, K.M, A.C. Manuscript writing by A.C and K.M. All authors contributed reviewing and editing the manuscript.

